# Biochat: a database for natural language processing of Gene Expression Omnibus data

**DOI:** 10.1101/480020

**Authors:** Bohdan B. Khomtchouk, Vsevolod Dyomkin, Kasra A. Vand, Themistocles Assimes, Or Gozani

## Abstract

A biological dataset’s metadata profile (e.g., study description, organism name, sequencing type, etc.) typically contains terse but descriptive textual information that can be used to link it with other similar biological datasets for the purpose of integrating omics data of different types to inform hypotheses and biological questions. Here we present Biochat, a database containing a multi-omics data integration support system to aid in cross-linking Gene Expression Omnibus (GEO) records to each other by metadata similarity through a user-friendly web application. Biochat is publicly available at: http://www.biochat.ai. Biochat source code is hosted at: https://github.com/Bohdan-Khomtchouk/Bio-chat.

**Database URL:** https://github.com/Bohdan-Khomtchouk/Bio-chat

## 1 Introduction

The Gene Expression Omnibus (GEO) is an international public functional genomics data repository supporting microarray, next-generation sequencing, and other forms of high-throughput functional genomic dataset submissions across a variety of gene expression studies that investigate a broad range of biological themes including disease, development, evolution, immunity, ecology, toxicology, metabolism, and other areas (1, 2, 3). In addition, GEO supports a variety of non-expression data representing diverse categories of functional genomic and epigenomic studies that include genome methylation, chromatin structure, copy number variations, and protein-DNA interactions submitted by the scientific community in compliance with grant or journal data sharing policies that require original research datasets to be made available in a public repository, the objective being to facilitate independent evaluation of results, re-analysis, and full access to all parts of a study (3). Since its inception, GEO has become one of the most popular database repositories for biomedical researchers to deposit their primary data as part of the research lifecycle. For example, a large proportion of primary research articles in PubMed often include a data availability section that lists a GEO accession identifier to access the array- or sequence-based data files associated with a study, including its associated metadata and other contents.

Given GEO’s penetrance across a wide spectrum of research fields and data types across the broader scientific community, understanding similarity between different studies (GSE accession identifiers) or datasets (GDS accession identifiers) may facilitate multi-omics integration. Since the biological data-verse is expanding every day, with new experimental data published daily, multidimensionally integrating this information at scale is essential to data-driven discovery. For example, bringing together ostensibly unrelated datasets (e.g., from different organisms, next-generation sequencing types, age groups, cell lines, etc.) can inform and contribute towards a deeper understanding of a variety of biological questions ranging from cancer to aging. To this end, quantifying what GEO records are most similar to any other given GEO record according to its textual metadata description (study description, organism name, sequencing type, etc.) would be useful for finding and computationally integrating GEO records along common themes or topics. One such possible workflow could be: read a paper → copy/paste its associated GEO accession identifier into a search bar → find other datasets/studies in GEO that have very similar metadata profiles, indicating actionable opportunities to explore those papers and their respective data, potentially integrating them in subsequent multi-omics follow-up studies to harbor biological insights that are not apparent when studying each dataset individually (4).

To date, GEO does not support such a functionality, which requires the development of natural language processing (NLP) algorithms trained on typically short and often sparse metadata fields. Although text mining of free text metadata has previously been shown to be promising for identifying related experiments through semantic similarity (5), bioNLP is still a largely underrepresented area in biomedical data science and constitutes an unmet need with respect to the development of multi-omics integration support systems.

NLP techniques have previously been used to design automated text mining methods that automatically identify disease-related experiments in GEO (6) and, more recently, NLP of text from GEO series was used to classify presence or absence of a disease signature, including classification of control vs. treatment samples based on metadata profiles (7). Additionally, tools to compare and contrast gene expression profiles based on automatic curation and NLP analysis of GEO records have also been developed (8). However, a multi-omics data integration support system for cross-linking GEO records by metadata similarity has not yet been devised. Therefore, given the variety of gene expression and non-expression data in GEO and the pace at which its growing -- doubling in size every 2 years, on average (9) -- we developed Biochat: an open-source publicly available web application for querying similar GEO records relative to each other using a variety of customized NLP algorithms and user-specified filters.

## 2 Methods

Biochat is implemented in Common Lisp and JavaScript running on an Amazon Elastic Compute Cloud (EC2) Linux instance. Similar to GEOmetadb (10), all datasets and studies within Biochat are faithfully parsed from GEO and no attempt is made to curate, semantically recode, or otherwise clean up GEO metadata field names. Due to the compute-intensive nature of algorithmically calculating a single GEO record’s similarity relative to the descriptive metadata profiles of over 100,000 existing GEO series (GSE) records, each search query takes (on average) between 2-3 minutes. Plans and ongoing work to use GPUs to pre-compute and cache similarity matrices on persistent storage (Amazon S3) to scale performance are currently underway (see Acknowledgements).

## 3 Results and Discussion

We developed Biochat, which is a collection of machine learning and natural language processing algorithms to group data records by similarity based on their free text description and other metadata information. The records are obtained from the datasets of biological experiments stored in the Gene Expression Omnibus (GEO) and automatically synced to be up-to-date with GEO every 24 hours.

Similarity, in the context of dataset descriptions, is not a well-defined concept, as the dataset record contains a number of metadata fields, including free-form text descriptive ones such as the title, summary, and experiment design, as well as more structured fields like the sample organism or sequencing platform. Besides, from the point of view of a researcher, different notions of similarity may be relevant. For instance, sometimes only datasets for a particular group of organisms are of interest. A more nuanced case is when only experiments that target a particular epigenetic factor (which may be mentioned in the text summary but is not indicated in a special field) are requested. That is why the Biochat project aims to provide a flexible toolset suitable for experimenting with different similarity measures and their parameters, as well as supplemental filtering based on additional settings.

In the context of Biochat, a similarity measure is a function of two records that returns a number in the interval [0,1] signifying the degree of similarity (the closer to 1 – the more similar). The magnitude of the similarity does not have any particular meaning, the only requirement is that records considered more similar should have a larger value of similarity. So, similarity values obtained by different similarity measures cannot be compared. Since different subdomains in biomedical literature vary along many linguistic dimensions, text mining systems performing well on one subdomain are not guaranteed to perform well on another (11, 12). Therefore, Biochat’s recommended usage is to try multiple different similarity measures one by one in the application’s user interface (UI) and examine the output results generated from each query -- ultimately using domain-specific expertise to compare and contrast the search results. Presumably, clicks on PubMed ID (PMID) hyperlinks in the UI (Figure 1) signify a level of active interest in learning more information beyond the content provided by the metadata fields alone.

**Fig. 1.**
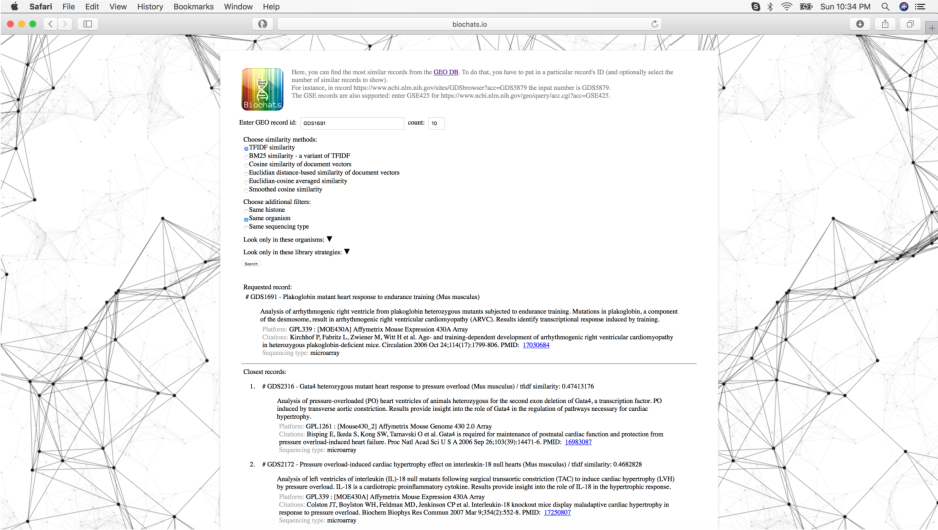
Biochat user interface. Searching for GDS1691 (Plakoglobin mutant heart response to endurance training) with TF-IDF similarity algorithm and same organism filter enabled returns GDS2316 (Gata4 heterozygous heart response to pressure overload) and GDS2172 (Pressure overload-induced cardiac hypertrophy effect on interleukin-18 null hearts) as the two closest records. Toggling different similarity methods and re-running the search each time is the recommended usage of this software program, as different algorithms often produce very different output results depending on the nature of the metadata.

In a long-term effort to leverage these clicks as a constant stream of collective intelligence that is gathered from a diverse community of users, we have pre-emptively implemented a PostgreSQL storage backend that captures user analytics to improve the Biochat search platform over time. Recorded information includes: IP address, timestamp, input query (GEO accession ID), output clicked on, and any NLP settings or user-specified filters. Our hypothesis is that real-time event stream processing of user clicks on PubMed ID (PMID) hyperlinks in the UI (Figure 1) represents a human-computer interface improvement that will ultimately refine the machine-based NLP approaches with human touch -- allowing us to learn patterns of biological data similarity above and beyond that provided solely by NLP algorithms. Over the years, we plan to evaluate this collective intelligence hypothesis in depth and release a follow-up paper reporting these results to the public, including results summarizing, e.g. what types of NLP algorithms perform most in accord with curiosity-driven domain expert clicks, ultimately testing the utility and efficacy of different algorithmic approaches across various domains of biological data science (Future directions).

Currently, in Biochat, two principle NLP approaches to similarity measurements are:

- bag-of-words-based similarity
- distributed representation-based similarity

In the bag-of-words (or token-based) approach, each record’s textual description is transformed into a sparse vector of the size equal to the size of the vocabulary. The transformation is performed by tokenization of the text, and then assigning some weight value to the element of the document vector representing each token. The bag-of-words similarity measures include the variants of term frequency-inverse document frequency (TF-IDF): vanilla one and BM25.

The TF and IDF vocabularies are calculated from the whole record set using the tokens from the record’s title, summary, and design description. TF count for each document is calculated as a ratio of token count by the document’s length. IDF count is calculated using the standard log weighting:

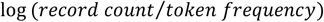

Both TF-IDF and BM25 similarity measure calculations use the stored weights. The similarity value of two records is calculated as a ratio of the sum of all TF-IDF weights of the tokens present in both record’s text descriptions divided by the product of the L2-norms of the TF-IDF vectors of each record.

The difference between the measures is that, in BM25, instead of the plain TF-IDF, the following formula is used:

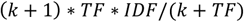

where *k* is the BM25 parameter, the default value of which is chosen to be 1.2.

Another approach to document representation implemented in Biochat is based on vector space models that use dense low-dimensional (vector size: 100-300) word vectors and combine them in some way into a same dimensional document vector. There are two approaches to obtaining the document vectors: by simple aggregation of the pre-calculated word vectors and by constructing the vector using a machine learning algorithm – see Paragraph vectors (13) or Skip Thought vectors (14). In Biochat, we chose to implement the aggregation approach using the PubMed word vectors (15) calculated with the word2vec algorithm (16). This is due to the availability of high-quality pre-trained vectors, lack of training data for the successful application of the doc2vec approaches, and the empirical results showing that simple word vectors aggregation performs not worse on short texts (17).

The similarity measures based on document vectors implemented in Biochat perform the comparison using the following algorithms:

- Cosine similarity and smoothed cosine similarity, where 5 is chosen as the default smoothing factor.
- Euclidian distance-based similarity. The formula for calculating the similarity score in the interval [0,1] is the following: 1/(*nrm2(v1 – v2)* + 1), where *nrm2* is the L2-norm.
- Combined cosine/Euclidian distance similarity that uses the square root of the product of both measures.

Since the main application of Biochat is sorting the GEO records database according to the similarity to a selected record, the sorted output may additionally be filtered based on the following set of criteria:

- Retain only records for a selected organism or group of organisms.
- Retain only records that mention a particular histone (e.g., H3K9me1) in its free text description.

## 4 Future directions

Since the biological research community is extremely diverse (e.g., a fly geneticist is likely to have a very different set of expertise/knowledge than an infectious disease immunologist or a bacteriologist), bringing these users to a common platform like Biochat could lead to a gradual paradigm shift in exploring biological data science. Specifically, observing and learning from the user dynamics of diverse domain experts interacting with metadata from a popular biological database like GEO over time may lead to a potentially valuable training set that could be used to refine the current NLP approaches developed in this paper. In general, computing on metadata descriptions through a seamless combination of human-based and machine learning-based (NLP) approaches may be a better and more effective strategy for finding emergent structure within large volumes of biological data -- and we plan to test this hypothesis further as Biochat accumulates more users over time. Therefore, the future direction of this work is to perform a hybridized study that is both data-driven and hypothesis-driven at the same time.

## 5 Acknowledgements

Some of the computing for this project is being performed on the Stanford Sherlock cluster. We would like to thank Stanford University and the Stanford Research Computing Center for providing computational resources and support that will contribute to the future directions of these research results. The authors declare no competing financial interests.

